# Portable and Error-Free DNA-Based Data Storage

**DOI:** 10.1101/079442

**Authors:** S. M. Hossein Tabatabaei Yazdi, Ryan Gabrys, Olgica Milenkovic

## Abstract

DNA-based data storage is an emerging nonvolatile memory technology of potentially unprecedented density, durability, and replication efficiency^1,2,3^,^4,5,6^. The basic system implementation steps include synthesizing DNA strings that contain user information and subsequently reading them via high-throughput sequencing technologies. All existing architectures enable reading and writing, while some also allow for editing^3^ and elementary sequencing error correction^3,4^. However, none of the current architectures offers error-free and random-access readouts from a portable device. Here we show through experimental and theoretical verification that such a platform may be easily implemented in practice using MinION sequencers. The gist of the approach is to design an integrated pipeline that encodes data to avoid synthesis and sequencing errors, enables random access through addressing, and leverages efficient portable nanopore sequencing via new anchored iterative alignment and insertion/deletion error-correcting codes. Our work represents the only known random access DNA-based data storage system that uses error-prone MinION sequencers and produces error-free readouts with the highest reported information rate and density.

---

Modern data storage systems primarily rely on optical and magnetic media to record massive volumes of data that may be efficiently accessed, retrieved, and copied^7^. Recently, these systems were challenged by the emergence of the first DNA- and polymer-based data storage platforms^1,2,3^,^4,5,8^. The proposed platforms have the potential to overcome existing bottlenecks of classical recorders as they offer ultrahigh storage densities on the order of 10^15^-10^20^ bytes per gram of DNA^1,2,3^,^5^. Experiments have shown that with DNA-based data storage technologies one can record files as large as 200 MB^5^, and ensure long-term data integrity through encapsulation^4^ and coding^5,9,10^. In addition, one can accommodate random access and rewriting features through specialized addressing^3^.

Current DNA-based data storage architectures rely on synthesizing the DNA content once and retrieving it many times. Retrieval is exclusively performed via high-throughput sequencing technologies, such as Illumina HiSeq^1,2^ and Illumina MiSeq^4,5^. These devices are designed for laboratories and are not portable. Portable sequencers exclusively use nanopores, which have been known to introduce a prohibitively large number of deletion, insertion, and substitution errors (with some estimates^11^ as high as 38%). At the same time, the error rates of modern magnetic and optical systems rarely exceed 1 bit in 10 terabytes^12^. Such high error rates have hindered the practical use of nanopore sequencers for DNA-based data storage applications^13^.

The two contributions of this work are a theoretical analysis and a practical implementation of a DNA-based data storage architecture that integrates error-control and constrained codes, sophisticated data post-processing techniques, and *portable, nanopore-based sequencing*. The data processing pipeline includes an encoding step performed before DNA synthesis and a post-processing step performed after sequencing.

## 1. The Encoding Step.

To accommodate large file sizes at low synthesis cost, the data is first compressed. Decompression may cause catastrophic error propagation if mismatches are introduced in the DNA strings either during synthesis or sequencing. Even one single substitution error in the compressed domain may render the file unrecognizable. To introduce the redundancy needed for different stages of error correction and minimize the addressing overhead, we chose the DNA codeword length to be 1,000 base pairs (bp). This codeword length also offers good assembly quality of long files without additional coverage redundancy or addressing, and the overall smallest commercial synthesis cost. To accommodate that choice of codeword length, as well as the inclusion of codeword address sequences, we grouped 123 × 14 = 1,722 consecutive bits in the compressed file and translated them into DNA blocks comprising 123 × 8 = 984 bases. We then balanced the GC-content of each substring of 8 bases via specialized constrained coding techniques that extend our previous results^14^, outlined in the Supplementary Information. Balancing eliminates some secondary structures, reduces synthesis errors, and helps to correct sequencing deletion errors. Each of the remaining 16 bases in a DNA codeword were used as codeword addresses. The purpose of the addressing method is to enable random access to codewords via highly selective PCR reactions. Selectivity is achieved by prohibiting the appearance of the address sequence anywhere in the encoded DNA blocks^3,14^. Additional protection against deletion errors is provided via *homopolymer check equations*. When coupled with balancing and subsequent read alignment steps, homopolymer checks lead to error-free readouts. A detailed description of the balancing and addressing schemes may be found in the Supplementary Information. Homopolymer checks are discussed in the post-processing step. All the encoding techniques are universal and therefore transparent to the type of data to be stored. They are illustrated in Figure 1.

**Figure 1.**
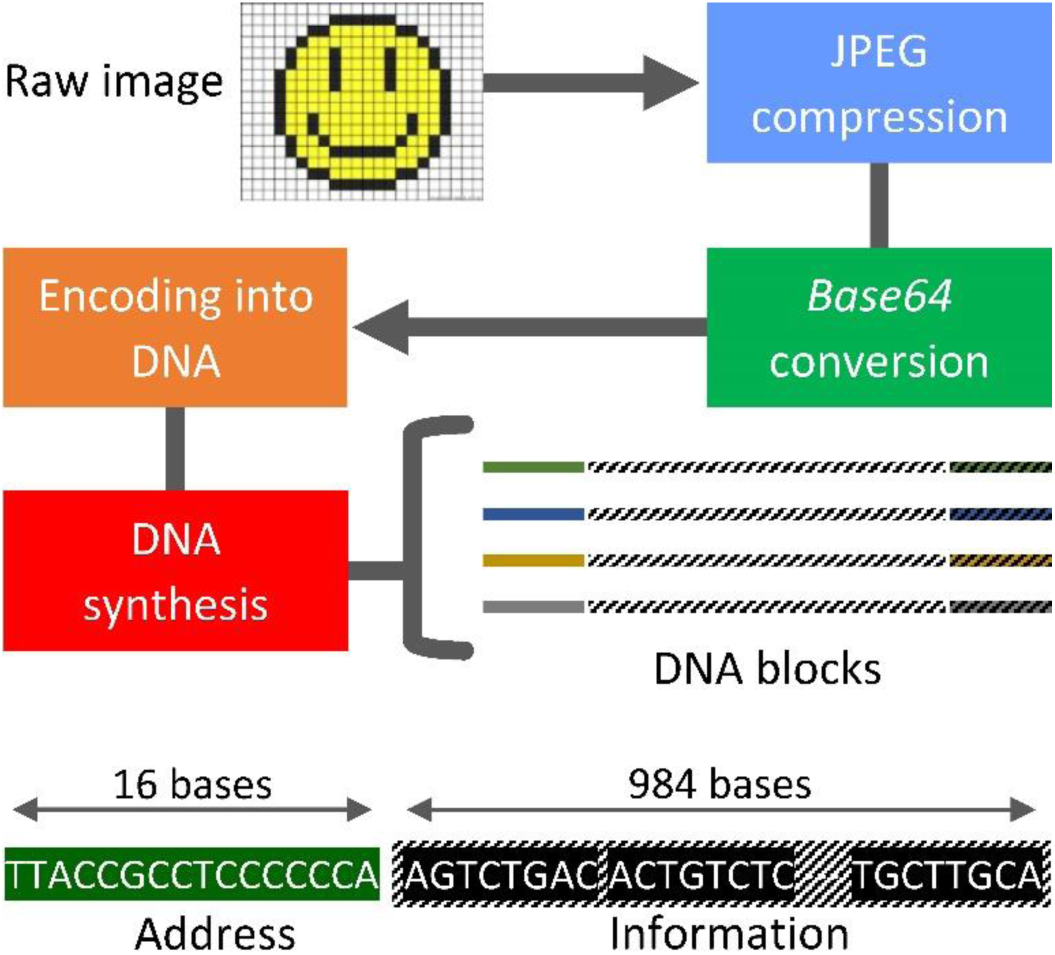
The encoding stage. This stage involves compression, representation conversion, encoding into DNA, and subsequent synthesis. Each synthesized DNA codeword is equipped with one or two addresses. The encoding phase entails constrained coding, which limits the occurrence of the address block to one predefined position in the codeword only, and GC-content balancing of each substring of eight bases. Additional homopolymer checks are added directly into the string or stored on classical media; they correspond to only 0.02% of the data content.

## 2. The Post-processing Step.

Our extensive testing reveals that reads obtained using the latest prototypes of MinION sequencers have sequence-dependent substitution, deletion, and insertion errors. In practice, arbitrary combinations of deletions, insertions and substitution are harder to correct than deletions alone. Hence, we performed a consensus alignment procedure that “transforms” almost all insertion and substitution errors to deletion errors confined to homopolymers of certain lengths, and generates an estimate of the DNA codeword based on the noisy reads.

In the first phase of post processing, we constructed a rough estimate of the DNA codewords. For that purpose, we used the address sequences to identify high-quality reads, i.e., those reads that contain an exact match with the given address. Aligning all reads instead of only high quality reads results in a large number of errors, and is to be avoided. Next, we ran different multiple sequence alignment (MSA) algorithms on the identified high-quality reads and obtained different consensus sequences. For that purpose, we used Kalign, Clustal Omega, Coffee, and MUSCLE^15,16^. As multiple sequence alignment algorithms are traditionally designed for phylogenetic analysis, their parameters are inappropriate for modeling “mutations” introduced by nanopore sequencers. Hence, for each alignment method, new parameters were chosen by trial and error (see the Supplementary Information). The choice of the parameters was governed by the edit distance between the MSA consensus sequence and the corresponding DNA codeword.

As each alignment method produced a different consensus sequence, we formed an aggregate consensus. The aggregate consensus contains the “majority homopolymer” of the different MSA algorithms. As an example, if three MSA algorithms produced three consensus sequences, AAATTGCC, AATTTGCA, and AAATTGC, the majority homopolymer consensus would equal AAATTGCA, as two sequences contain a homopolymer of three As at the first position; two sequences contain a homopolymer of two Ts in the positions to follow; and all three sequences contain G and C. Observe that A is included in the last position of the consensus.

In the second phase of post processing, we performed iterative alignment. By this stage, consensus sequences that estimate the original DNA blocks were identified, with errors mostly confined to deletions in homopolymers of length at least two. (See the Supplementary Information). To further improve the reconstruction quality of the blocks and thereby correct more errors, we performed one more round of BWA^17^ alignment to match more reads with the corresponding estimates of their DNA codewords. Once this alignment was generated, two sequential checks were performed simultaneously on the bases. The checks included computing the majority consensus for each homopolymer length and determining whether the GC-balancing constraint for all substrings of length 8 was satisfied. More precisely, in the majority count, only homopolymer lengths that resulted in a correct balance were considered. This procedure is illustrated by an example in the Supplementary Information. Note that alignment does not require any coding redundancy, while balancing uses *typical sequences* and, as a result of this, has a high coding rate of 0.88. The alignment procedure is depicted in Figure 2.

**Figure 2.**
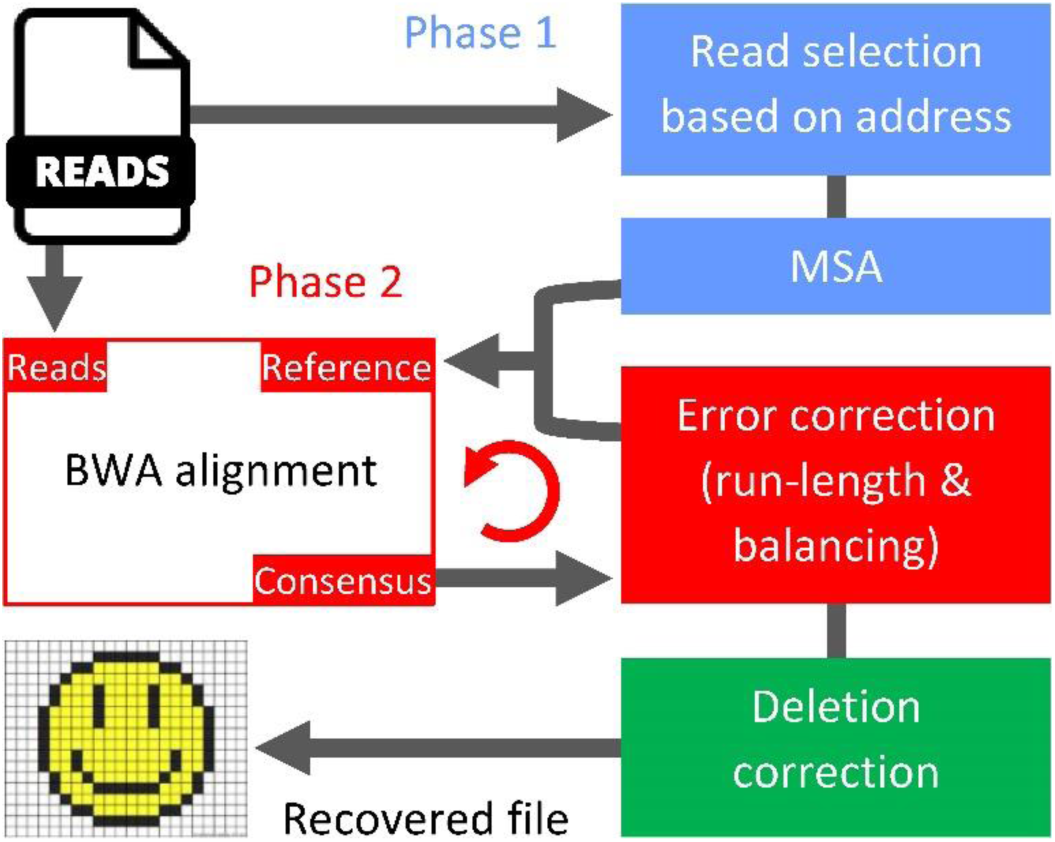
Post processing via sequence alignment and homopolymer correction. In the first phase, estimates of the DNA codewords are obtained by running several MSA algorithms on high-quality reads that contain an exact match with the address sequence. The second phase improves the estimate by employing an iterative method that includes BWA alignment and an error-correcting scheme.

In the final stage of post processing, we corrected deletion errors in homopolymers of length exceeding one. For this purpose, we used an error-correction scheme that parses the consensus sequence into homopolymers. As an example, the parsing of the sequence AATCCCGA into homopolymers AA, T, CCC, G, A gives rise to a homopolymer length sequence of 2,1,3,1,1. Special redundancy that protects against asymmetric substitution errors is incorporated into the homopolymer length sequence.

If two deletions were to occur in the example consensus, resulting in ATCCGA, the homopolymer lengths would equal 1,1,2,1,1. Here, we can recover the original length sequence 2,1,3,1,1 from 1,1,2,1,1 by correcting two bounded magnitude substitution errors. Note that the sequence of the homopolymer symbols is known from the consensus.

Because we extensively tested address-based DNA data storage methods for ordinary text files^3^, for practical implementation we focused on image data. Two images were used as samples: A poster for the movie Citizen Kane (released in 1941), and a color Smiley Face emoji. The total size of the images was 10,894 bytes. The two images were compressed into a JPEG^18^ format and then converted into a binary string using *Base64*^19^ (*Base64* allows one to embed images into HTML files). The resulting size for the two compressed images was 3,633 bytes.

Through the previously described data encoding methods, the images were converted into 17 DNA blocks, out of which 16 blocks were of length 1,000 bp and one single block was of length 880 bp. Before the sequences were submitted for synthesis, they were tested by the IDT (Integrated DNA Technologies) gBlocks® Gene Fragments Entry online software; they were then synthesized. The total cost of the testing and synthesis was $2,540. IDT failed to synthesize one of the blocks because of a high GC-content in one substring of the address sequence, which was subsequently corrected through the addition of adapters at the two ends of the sequences. Based on information about this type of synthesis error, the sequence encoding procedure was modified to accommodate balancing of all short substrings of the DNA blocks, including the addresses, as previously described.

The gBlocks representing our DNA codewords synthesized by IDT were mixed in equimolar concentration. One microgram of pooled gBlocks was used to construct the Oxford Nanopore libraries with the Nanopore Sequencing kit SQK-MAP006. The gBlock libraries were pooled and sequenced for 48 hours in a *portable size* MinION MkI using a flowcell Mk 1 FLO-MAP103.12. All of the reads used in our subsequent testing were generated within the first 12 hours of sequencing. Base-calling was performed in real time with the cloud service of Metrichor (Oxford, UK); the run produced a total of 6,660 reads that passed the filter. The sequencing error rate was 12%, a significant improvement over previous architectures^11^. After the consensus formation stage, the error rate reduced to a mere 0.02% without any error-correction redundancy. The three errors in the 17 codewords were subsequently corrected using homopolymer checks, thereby producing error-free reconstructed images. The images reconstructed with and without homopolymer checks are shown in Figure 3.

**Figure 3.**
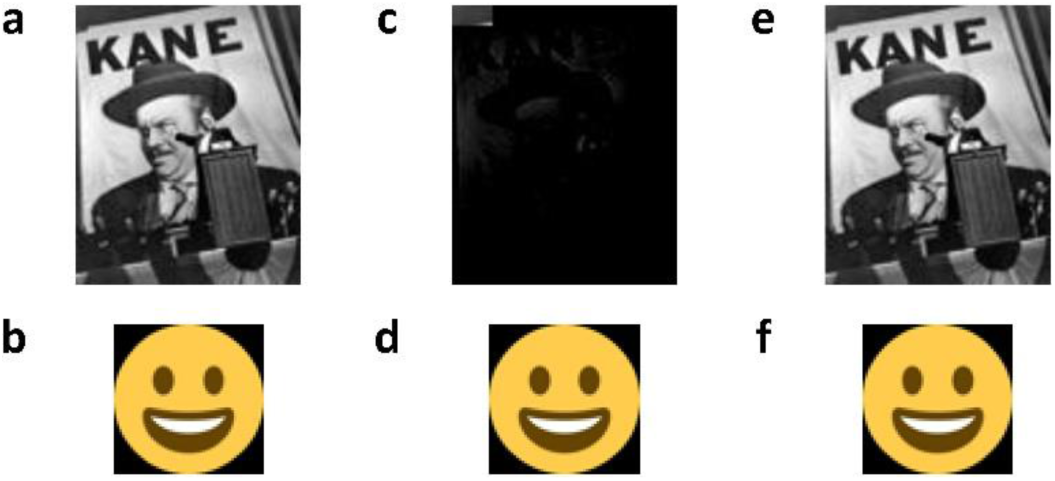
Image files used in our experiment. **a, b,** show the raw images which were compressed, encoded and synthesized into DNA blocks. The Citizen Kane poster and Smiley Face emoji were of size 9,592 and 130.2 bytes, and had dimensions of 88×109 and 56×56 pixels, respectively. **c, d,** show the recovered images after sequencing of the DNA blocks and the post-processing phase without homopolymer error correction. Despite having only two errors in the Citizen Kane file, we were not able to recover any detail in the image. On the other hand, one error in the Smiley Face emoji did not cause any visible distortion. **e, f,** show the image files obtained after homopolymer error correction, leading to an error-free reconstruction of the original file.

The described implementation represents the only known random access DNA storage system that operates in conjunction with a MinION sequencer. Despite the fact that MinION has significantly higher error rates than Illumina sequencers and that random access DNA systems typically require additional data redundancy, our DNA storage system has the highest reported information rate of 0.85, storage density of 1.1 × 10^23^ bytes/gram, and it offers error-free reconstruction.

## Acknowledgments

We thank Alvaro Hernandez for running the MinION experiments and for many valuable discussions, and Jenny Applequist and Chrissy Gabrys for providing feedback on the manuscript. We also gratefully acknowledge funding under the NSF grants CCF 1618366 and NCF CSoI Class 2010 grant.

## Author Contributions

O. M. conceived the research. S. Y. performed the algorithmic implementations for alignment. R. G. developed the homopolymer codes. S. Y, R. G. and O. M. developed the post-processing scheme and wrote the paper.

